# High frequency monitoring of feeding activity in benthic suspension feeders

**DOI:** 10.1101/2022.11.29.518401

**Authors:** Antonio Agüera, Tore Strohmeier, Cathinka Krogness, Øivind Strand

## Abstract

Suspension feeders ecosystem role and services are mainly driven by their efficiency in clearing particles from the water column. As such there is an interest on suspension feeders feeding activity and how they interact with the ecosystem. Advancing research on feeding response requires experimental designs where individuals can be exposed to continuously changing environments and where feeding activity can be monitored at a high frequency and at different time scales. There are several methods to monitor feeding activity (as clearance rate (CR) and particle capture efficiency (CE)) of suspension feeders. Among them, the flow-through method is most often used and it allows individuals to be exposed to changing conditions. However, as with the other existing methods, the flow-through method is labor intensive and time consuming limiting both the number of individuals that could be observed simultaneously and the resolution at which measurements can be taken. The flow-through method is constrained by the need to measure the flow rate through the chambers and to take samples to determine particle concentration in the water. In this work we automated the standard flow-through method using micro-controller based prototyping. The result is a methodological approach that continuously monitor feeding at high frequency and on a larger number of individuals while reducing handling and measurement errors. This work provides the description and assessment of the automated set-up, which is an end to end solution that can readily assembled and configured.

## 1. INTRODUCTION

Benthic suspension feeders play an important ecological role in aquatic ecosystems [1]. They provide a vast variety of ecosystem services, going from improving water quality to food production. Their feeding activity is the main driver of these ecosystem services [2] and backs the research effort on suspension feeders feeding physiology. Nowadays, there is a growing interest in how environmental conditions affect suspension feeders feeding activity and how suspension feeders impact ecosystems [3, 2, 4, 5]. The determination of feeding (i.e. particle clearance rate and capture, transport, sorting and ingestion of suspended particles) is the core task of research studies on suspension feeders physiology [6, 7, 5, 8, 9]. There are several methods to determine these traits of suspension feeding: flow-through, closed system, biodeposition, suction, etc. These methods have different implementation, yet all of them are labor and time intensive and require supervision. This constrains experimental designs limiting the the number of replicates and the frequency of measurements [8]. Advancing research on feeding physiology requires experiments designed to observe suspension feeders under natural conditions or under combinations of stressors that change over time [10, 11]. The time scale at which these factors may affect the physiological response [10] have prompted the need to increase the frequency of measurements both across and within individuals [5].

Among the existing methods, the flow-through feeding method is widely used to measure clearance rate (CR) and capture efficiency (CE) of suspension feeders [7, 3]. This method consists of an open circulation system where individuals are exposed to running water. This flow through chamber design allows for individuals to be continuously exposed to varying environments and treatments with minimal handling [12, 10, 11, 4, 5]. In the flow-through method feeding is determined by measuring the depletion of suspended particles within the chamber at a measured flow rate. Flow rates need to be measured accurately and be within an optimal range referred as the flow independency phase [8]. Flow rates outside the flow independency phase result in erroneous determination of CR [8]. Flow rates are measured manually using a graduated cylinder and a timing device providing an unknown and variable error related to observer and measuring conditions. Depletion of suspended particles is calculated by counting and/or determining the size distribution of suspended particles on water samples from the outflow of the feeding and control chambers using flow cytometry [13], an electric [12] or a laser particle counter [4]. The current practice to determine CE and CR using the flow-through method is a time intensive manual process and thereby imposes a strict trade-off between the number of individuals observed simultaneously and the replication of measurements in time.

The use of microcontrollers and prototyping tools to undertake laboratory and field monitoring tasks offer a wide range of options to design experimental set-ups. Among other applications micro-controllers allow for automation and handling of data recording, prototyping of mechanical parts and sensors and interfacing with other instruments [14, 15, 5]. Microcontroller prototyping can be use to build systems to execute manual and supervised tasks automatically and/or remotely [16]. This work describes a microcontroller based prototype that builds on the design and assumptions of the flow-through method and automate the time consuming tasks involved in the standard methodology. The automated set-up monitors and records flow rates and collects and analyses water samples in the flow-through method. The final result is a method that continuously sample feeding activity at high frequency over long periods of time and that has the capacity to observe more individuals simultaneously while reducing individual manipulation and observer errors.

## 2. MATERIAL AND PROCEDURES

The method set-up consists of a rack of 22 feeding chambers designed for the application of the flow-through method with *Mytilus ssp* [12, 8]. This flow-through set-up is automated to increase the frequency, efficiency and accuracy of flow and suspended particles concentration (Figure 1). The setup is flexible and modular allowing for the adaptation to different sets of chambers to be used with different species or size classes.

**Figure 1:**
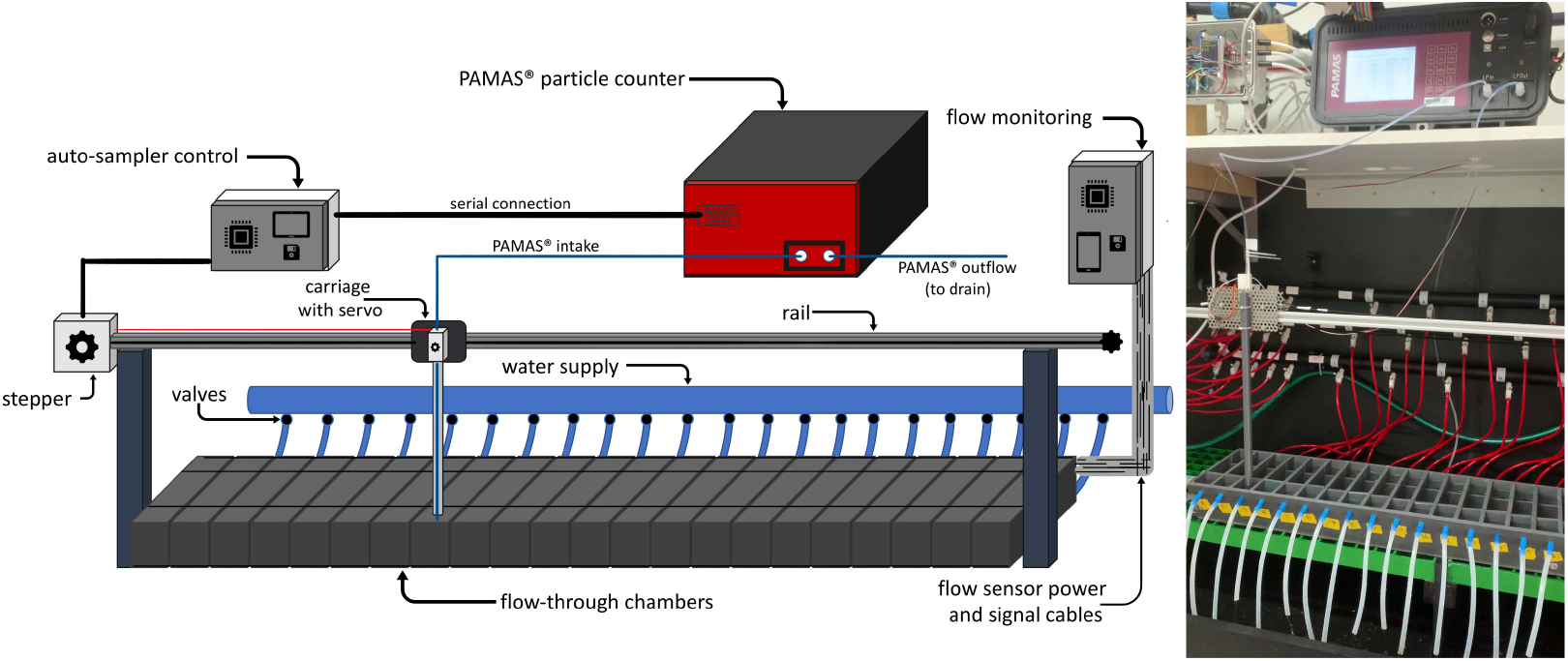
Automatic system set-up. Left: diagram showing the main components: PAMAS S4031GO, autosampler, control units (microprocessors) and chambers. Right: detail of built system working at the laboratory.

### 2.1. Automation of flow rate monitoring

Seawater is provided to the system in excess to maintain sufficient flow in the feeding chambers. Flow into each chamber was regulated using a needle valve that allowed setting a flow rate within 100-300 *ml min*^−1^. A flow sensor based on the Hall effect was fitted to the water line feeding each chamber (Figure 2). Hall effect flow sensors are based on paddles that turn as water pass through them. The sensor follows the movement of the paddles producing a signal that is used to count the turns. The frequency at which the paddles turn is correlated with the water flow-rate [17]. These sensors are widely used in industrial applications to keep track of dispensing volumes and flow rates, they are available in several configurations with variable accuracy, mounting constrains and working ranges from a few milliliters to several liters per minute [17, 18]. This set-up uses flow sensors manufactured by Shenzhen EPT technology Co. Ltd with a range from 0.1 to 3 *l min*^−1^ (Figure 2). Water supply, valves and flow sensors were selected to work within the necessary flow rate range for the animals used during the assessment of the method according to previous experience [12, 5].

**Figure 2:**
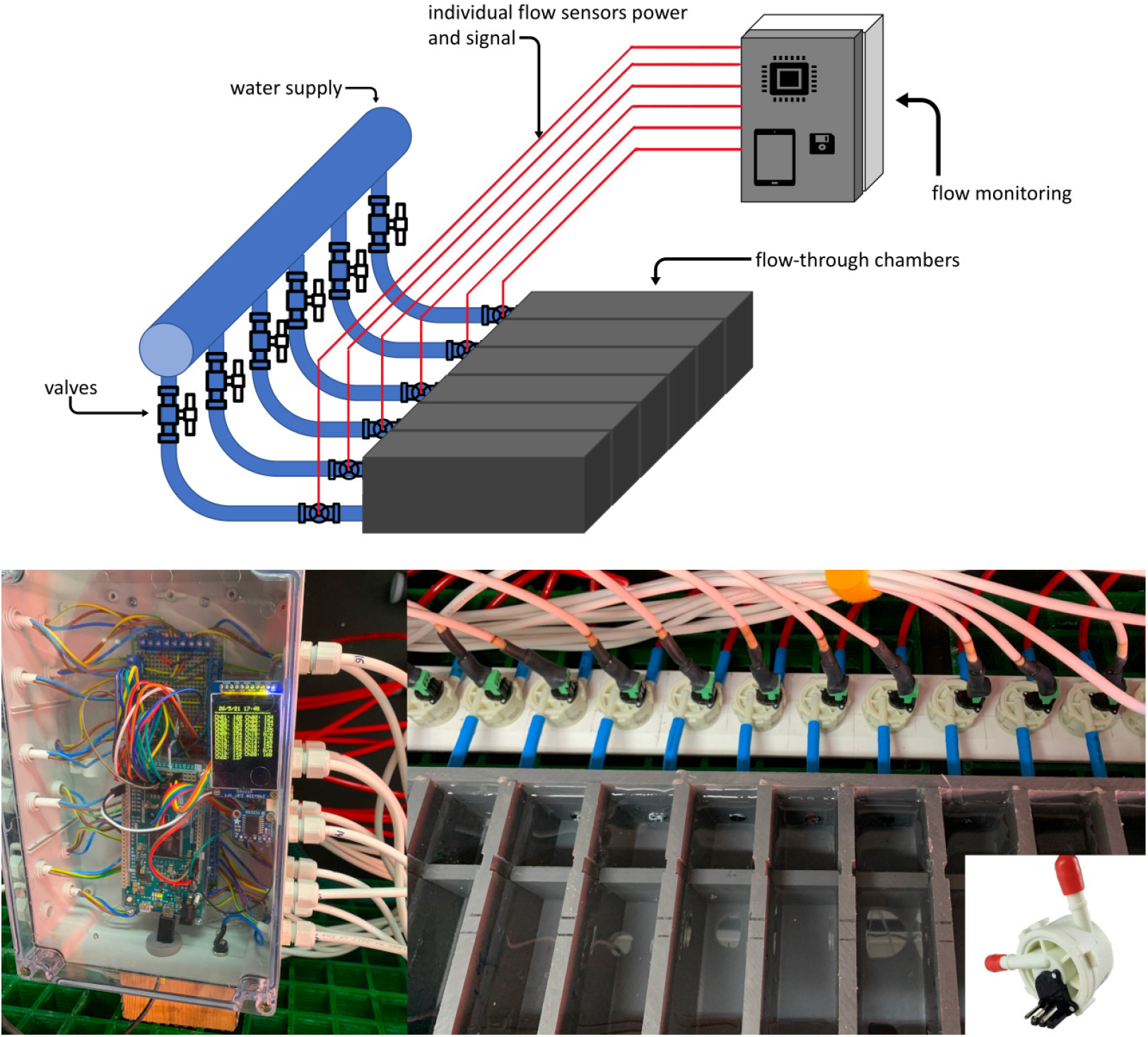
Automatic water flow monitoring system. Top: diagram showing wiring and arrangement of valves and flow sensors in the set-up. Bottom left: detail on the flow monitoring unit with microcontroller, tft screen and real time clock. Bottom right: Detail of flow sensors and chambers. Miniature detail of the EPT flow sensor used.

The signal from the flow sensors is monitored and recorded using an Arduino^®^ Mega 2560 rev3 microcontroller (https://docs.arduino.cc/). Each flow sensor output was attached to a different digital pin (22 pins were use in this case) which was configured as input pin and pulled-up [19]. The microcontroller was also fitted with a Adafruit DS3231 Real Time Clock breakout and 2” ST7789 TFT screen and with a micro-SD card slot (Adafruit Industries, LLC, New York, USA). This microcontroller was programmed to monitor the flow rate in the chambers. Flow rate of each chamber was monitored for 5 seconds every 2 minutes, the recorded values were timestamped and saved in a SD card and the last measurement in *ml min*^−1^ for each chamber was shown in the TFT screen (Figure 2). The flow sensors calibration constant (*turns ml*^−1^) were estimated passing a known volume of water through them while being monitored by the microcontroller (details of component wiring in Figure Appendix A.2).

### 2.2. Automation of water sampling and determination of suspended particles

The method uses a laser particle counter PAMAS S4031GO (PAMAS GmbH, Hamburg, Germany) to determine the concentration and size distribution of suspended particles in the water. This instrument has been used extensively in suspension feeder feeding physiology studies to understand CE, size selectivity and determine CR [4, 20, 21, 22]. The PAMAS uses light scattering to count particles by size class in predetermined intervals. The PAMAS incorporates its own pumping system and it registers calibrations, sampling profiles, samples ID and results. As a standalone system it only requires that the samples are taken to the intake line of the PAMAS. The PAMAS can be set to sample from the intake following a sampling profile programmed within the instrument, however the number of replicates is limited and it still needs the operator to supply the new sample when needed and input any sample identification. This constrains the upscaling of the set-up in number of replicates and measuring frequency. To solve the shortcomings of the PAMAS, another Arduino Mega 2560 rev3 was used to interface the PAMAS with an autosampler. PAMAS instruments are available with USB and serial (RS-232) interface and can be controlled and configured remotely using commands through serial communication.

The autosampler takes the intake line of the PAMAS to the desired chamber to be sampled and give the PAMAS the needed instructions (size range and configuration, sampling time, flushing time, etc). The autosampler maintain the PAMAS intake position while it is sampling and processing the sample (Figure 1). The PAMAS intake line was fixed to a carriage that moves along an aluminium rail over the feeding chambers. A bipolar stepper motor (12v, 350mA, NEMA-17 size) was used to control the position of the carriage using a timing belt. The carriage was fitted with a linear servo rail which moved the PAMAS intake line vertically to lower or lift it from the chambers when needed. Water sampling by the PAMAS was done by the chamber outlet. To control the motor and the servo the Arduino was fitted with a Adafruit Motor Shield v2.3 (Adafruit Industries, LLC, New York, USA (Figure Appendix A.1). The PAMAS was controlled from the same Arduino using a RS-232 to TTL converter (MAX3232IDR by SeeedStudio, Shenzhen, China). Serial communication with the PAMAS was done following the commands and instructions provided by the manufacturer. The PAMAS used in this work did not have a serial interface accessible (as a DB9 port) as standard. However, the manufacturer provided the needed information to access this interface on the main board of the instrument. The micro-controller was also fitted with an Adafruit DS3231 Real Time Clock breakout and ST7789 TFT screen with a micro-SD card slot.

The autosampler microcontroller was programmed to configure the PAMAS on start-up providing it with the calibration information, the size configuration of the channels to be used (a maximum of 32 channels that determine the predefined size ranges to measure) and the desired sampling profile (flushing time, sampling time/volume and number of replicates). Other parameters needed by the microcontroller are the number of chambers in the set-up, the distance between chambers and the number of the chambers to be used as controls. After configuring the PAMAS the microcontroller starts sampling the first chamber: the auto-sampler lowers the linear servo to the water and instruct the PAMAS to initiate the sampling profile. PAMAS sampling involves flushing the line during a time long enough to make sure the water from the current chamber has reached and filled the PAMAS system and counting particles in one or several samples of a predefined volume. After the instructed sampling is concluded the microcontroller requests the results from the PAMAS and stores them in the SD card timestamped, with the chamber number and the metadata information of the used sampling profile. Then the microcontroller lifts the line and moves the carriage to the next chamber to initiate the next sampling. Besides requesting the raw data from the PAMAS, the microcontroller can perform some calculations and data processing directly. As such, in some applications the program can use the information from the control chamber to calculate retention of a specific particle size range in all the chambers and display it at the screen. This feedback helps the operator to keep control of the experimental settings (flow rates, sampling volume, food treatments) and adjust them as needed. Recorded data can also be used by the system itself as a trigger or to control other instruments in the set-up. When the autosampler samples the last chamber of the rack the carriage is moved back to start over immediately or after a defined interval. Depending on the PAMAS sampling profile used, all 22 chambers could be sampled in less than 45 minutes.

Arduino IDE sketches along with auxiliary libraries and wiring diagrams can be found at https://git.imr.no/low-trophic-aquaculture/arduin-lab-codes/hf-flow-through.

## 3. ASSESSMENT

The method was assessed in an experimental trial at Øydvinstod, Ulvik, Norway in June 2021. The set-up was assembled in a laboratory with water supply from a pier next to it. Two pumps, delivered seawater from three meter depth into two header tanks. From the header tanks two diaphragm pumps with a pressure switch (Flojet RLF222201D) provided water to the chambers at a rate of 3.8 *l min*^−1^ and maximum pressure of 2.5 bars. Half of the chambers were connected to each water supply. Nineteen mussels (*Mytilus ssp*, 4.5-5 cm shell length) were collected from a culture net were placed in the feeding chambers. Two chambers (one with water from each supply) were left empty as controls for each supply. The aim of this trial was to continuously measure flow and CR for this population during five days at decreasing depth of one of the pumps set at sea. This set-up exposed the mussels to changes in environmental conditions during the experiment (Appendix B.1).

The chambers flow rates were set between 125 and 250 *ml min*^−1^ at the beginning of the experiment. The auto-sampler was set to sample continuously using 2 minutes per chamber (1 minute for flushing and another minute to sample 10 ml of seawater). This timing resulted in sampling all 21 chambers every 42 minutes. Results recorded by both microcontrollers were used to calculate CR following Steeves et al. [5]. At the end of the experiment each individual chamber was sampled 125 times (i.e. a total of 2375 CR measurements of 19 individuals). Flow rate through each chamber was measured over 1630 times.

The calibration of the 21 flow sensors provided a common calibration constant of 5.94±0.29 *turns ml*^−1^ (N = 63). The micro-controller can only count complete turns, meaning that the error in the count recorded was of at least ±1 turn (i.e. ±0.17ml). When measuring flow for 5 seconds this count error translated to a measuring error of 2% at 100*ml min*^−1^ going down to 0.6% at 300*ml min*^−1^. The results showed that the flow rate in some chambers drifted rapidly out of the targeted intervals. There was a static (open/closed) respirometry system connected to the same water supply, when the respiration system was closed the flow into the feeding chambers increased and the flow decreased every time part of the water was diverted to flush the respiration chambers. However even without readjusting flows, the flow rate range was maintained within the flow independency phase and below the flow inhibition phase most of the time providing appropriate data for the determination of CR (Figure 3).

**Figure 3:**
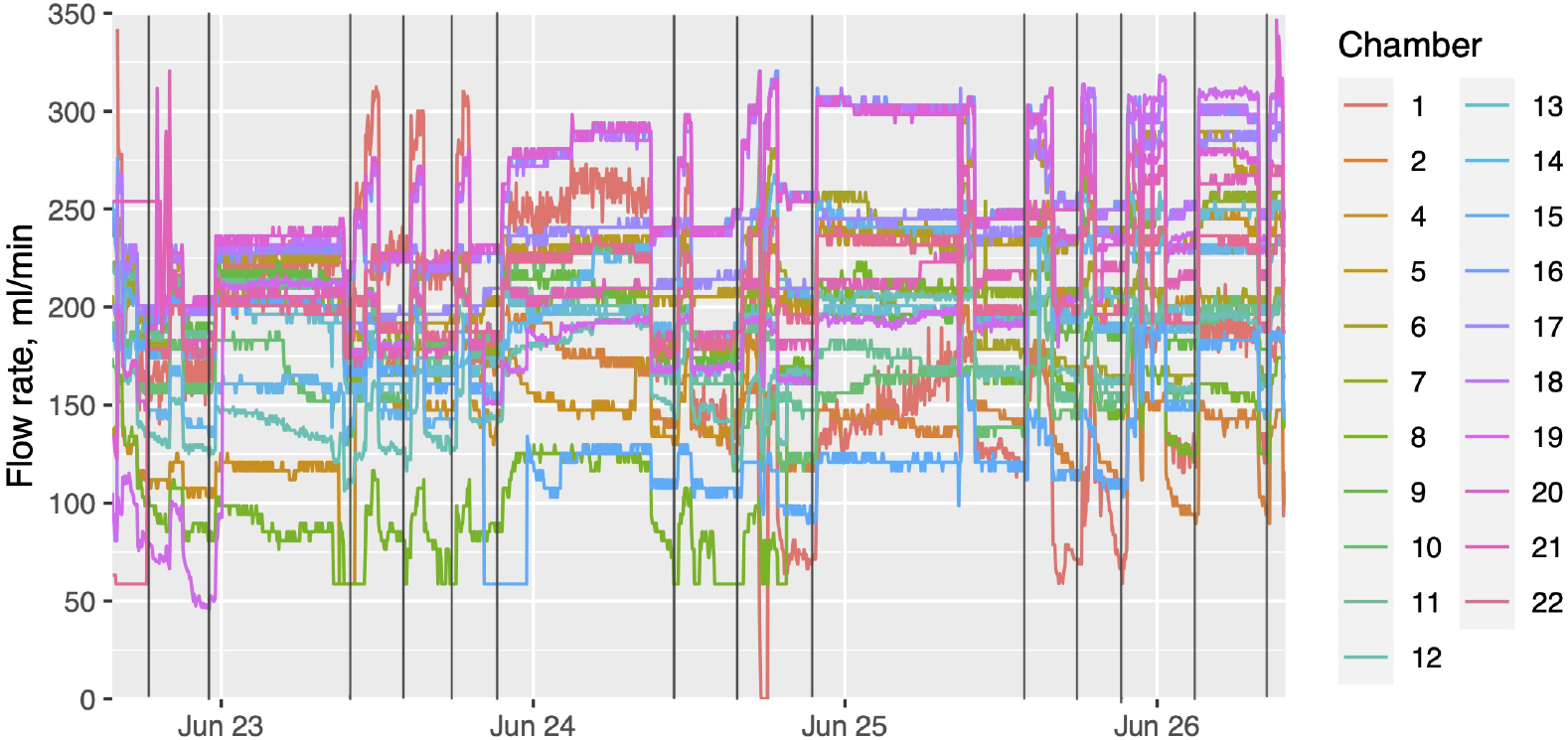
Measured flows in all 21 chambers during the trial. Black lines indicates when water to respirometry system was closed. Depending on the time of the day this lasted just a couple hours or all night.

The autosampler showed stable functionality during the experiment. Interfacing the PAMAS to the autosampler and controlling the processes of timing and synchronization with the micro-controller made it possible to increase the frequency of sampling and to collect data continuously. The microcontroller data handling avoided the need to name each sample in the PAMAS manually before processing or keeping track of multiple water samples from the different chambers. Timestamp on both microcontrollers records allowed for fast and accurate merging of the data. This ensured CR was determined at flow rates measured within ± 1 min. The flow rate variability observed during the experiment showed the effect of flow rate on the determined CR (Figure 4). The decrease of CR at flows below 100 - 120 *ml min*^−1^ suggest those measurements were taken within the flow dependency phase. Meanwhile the determination of CR at flows above 120 *ml min*^−1^ are within the flow independency phase and below flow rates within the feeding inhibition phase. These results agree with previous studies using the same chambers and similarly sized mussels feeding on natural seston [8].

**Figure 4:**
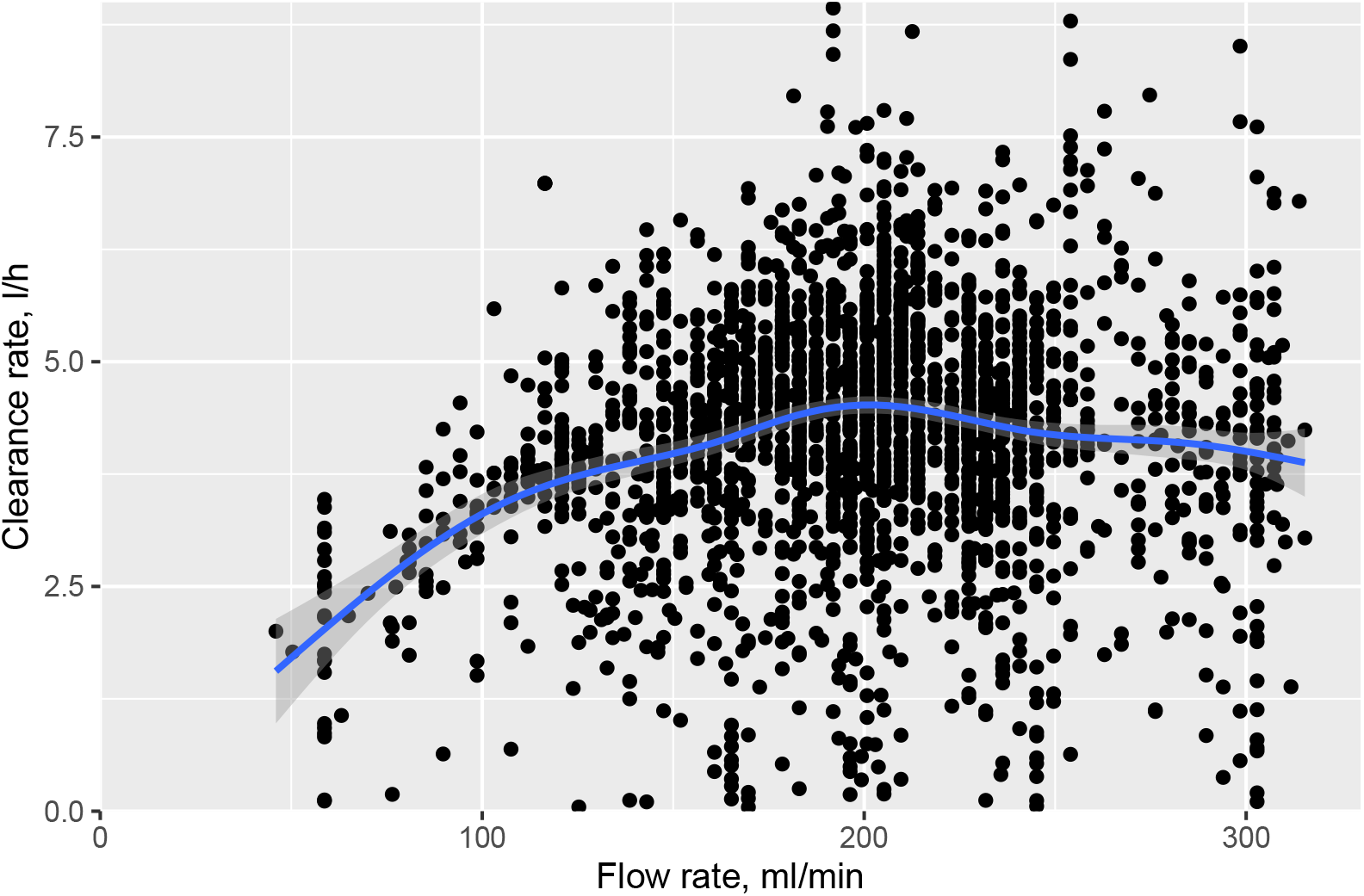
Calculated clearance rates of *Mytilus spp*. of 4.5-5 cm shell length at measured flow rates. Points are observations, blue line is a General Additive Model smoother, shaded area is model standard error.

Clearance rates calculated during the experiment were within rates measured for this population and mussels of similar size available in the literature [4, 12]. Variability in CR depends on many factors, including inter- and intraindividual variability and environmental conditions, making any statistical comparison with previously published data difficult. Direct observation of the data shows how the method is able to show differences between mussel under the different seawater sources and changes at the timescale at which environmental changes were observed (Figure 5, Figure Appendix B.1).

**Figure 5:**
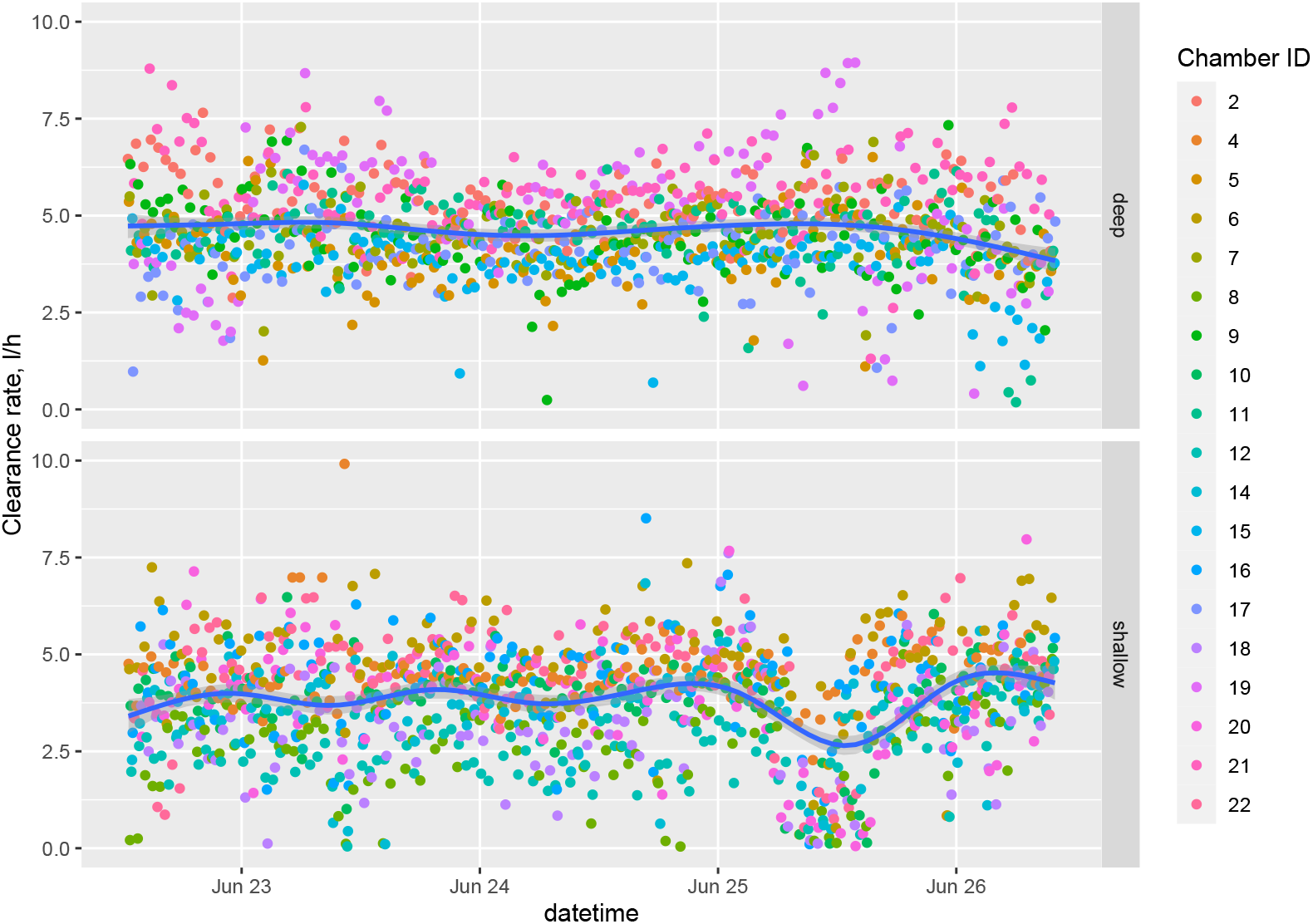
Individual clearance rates of *Mytilus spp*. of 4.5-5 cm shell length measured during the experiments. Treatments deep and shallow corresponded with the two pumps set at different depths. Shallow pump was raised daily forcing environmental changes in those individuals receiving water from that pump

## 4. DISCUSSION

Microcontroller prototyping offers a wide variety of solutions to automate and improve laboratory routine work and complex tasks [15, 16]. The microcontroller prototype presented here has taken a time and labor consuming laboratory methodology into a fully automated procedure. The resulting methodological approach is an end to end solution to advance research on suspension feeders ecology and physiology. The automated approach becomes a flow-through method that can be run continuously for several days providing a new sample each second minute with only one laser particle counter. The assessment trial showed how this method can be applied to gather high frequency data on the response of feeding activity of suspension feeders under a continuously changing environment. High frequency data means that the method have the capacity to observe short-term changes in the feeding response of suspension feeders under a continuously changing environment. Previous attempts to measure mean short term changes in CR in bivalves have obtained three to five samples a day [12]. In comparison, the assessment trial obtained 34 CR samples every day.

Using sensors to monitor individual chambers flow rates provided monitoring and control over the experimental conditions and ensures that correct flow rate is used for CR calculations. Flow rates outside the flow independency phase [8] (Figure 4 can also be detected and handled. This is more relevant in longer experiments where flow rates can change significantly over time due to several factors including tides, pumps effectiveness, changes in water properties, etc. Compared to the use of flow sensors, manual determination of flow rates is time consuming and results have a variable unknown error. How the sensors error compares with the error of measuring flow using a graduated cylinder and a timing device is difficult to assess as it depends on the cylinder precision, the reaction time of the observer and the flow rate being measurement.

The autosampler automated a fundamental task of the flow-through method, undertaking unsupervised continuous sampling of the feeding chambers. PA-MAS instruments are widely used and are important tools in studies on suspension feeders physiology [4, 20, 21, 22]. In this set-up the PAMAS possibilities are no longer restricted by the instrument software and the PAMAS becomes a more flexible and efficient tool that can be set-up to meet the experimental needs. As such, the system can sample the same chamber several times with different size intervals each time, or different sampling volume to assess sampling errors. The data is gathered and stored by the microcontrollers and the user can decide how this is handled or already program the microcontroller to process the data as needed before being stored. Further integration between the two microcontrollers used in this set-up is also possible. The application presented here benefited from high frequency recording flow rates. However it is possible for both microcontrollers to work together. For example, the autosampler microcontroller can be programmed to query the flow rate monitoring microcontroller to measure flow in a specific chamber for a given time and return that measurement back. As such this set-up provides an ample space to tailor the method to the experiment needs.

Monitoring feeding activity with this method can be used to better resolve the effect of feeding activity on the ecosystem services of suspension feeders. The automatic high frequency monitoring of suspension feeders allows to perform experiments to study short term feeding responses to fast changes in environmental conditions, such as food, temperature, salinity. With this approach the monitoring of feeding processes can be done at the relevant frequency to assess the plasticity at the individual and population levels. This automatic approach allows to upscale the set-up to accommodate a large number of individuals which increases the flexibility of the experimental design.

## Appendix A. Microcontrollers units and wiring

**Figure Appendix A.1:**
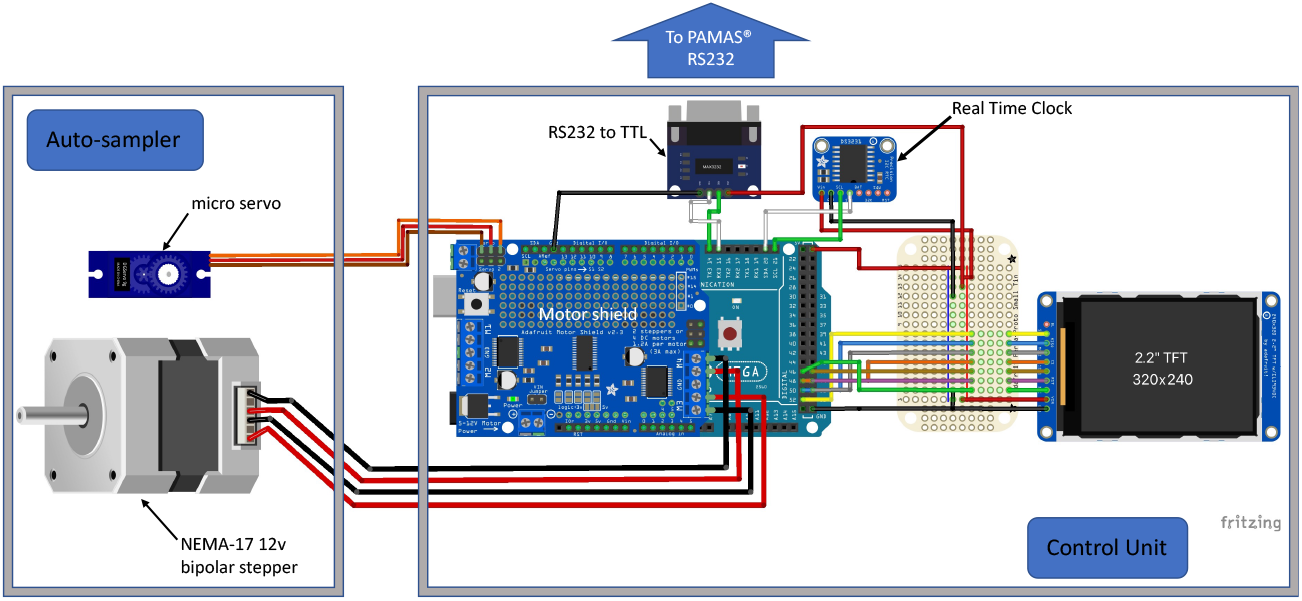
Schematic drawing showing the interfacing and wiring of the components used to control the laser particle counter and the autosampler.

**Figure Appendix A.2:**
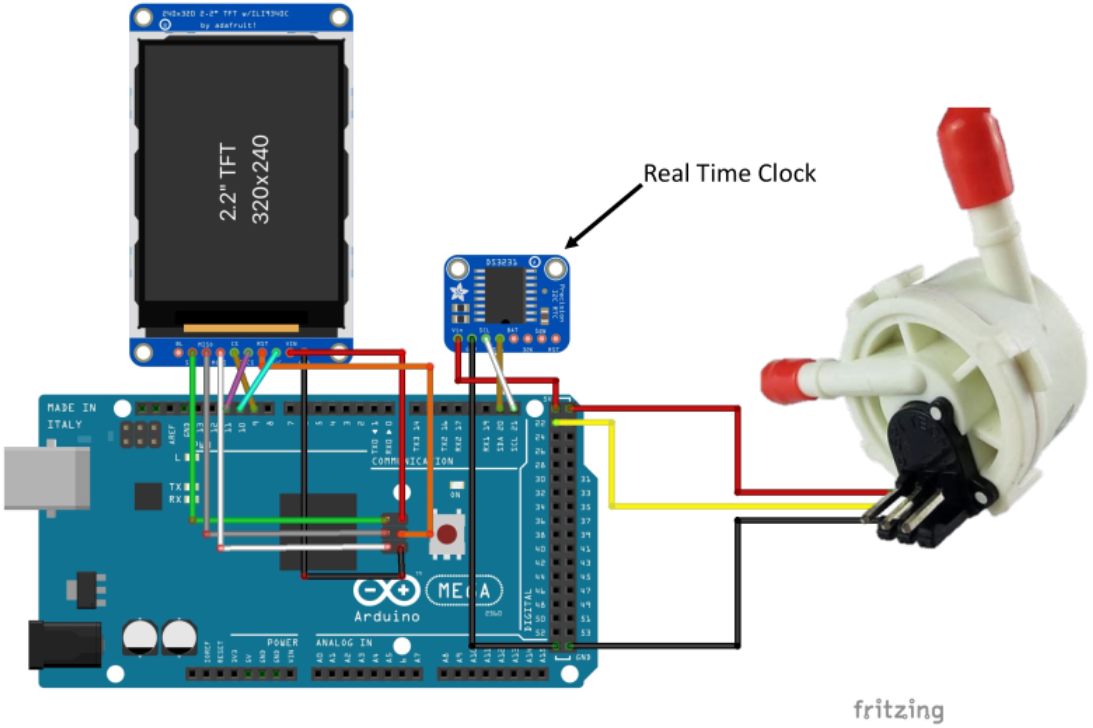
Schematic drawing showing the interfacing and wiring of the components of the flow rate monitoring system

## Appendix B. Environmental conditions during the experimental trial

**Figure Appendix B.1:**
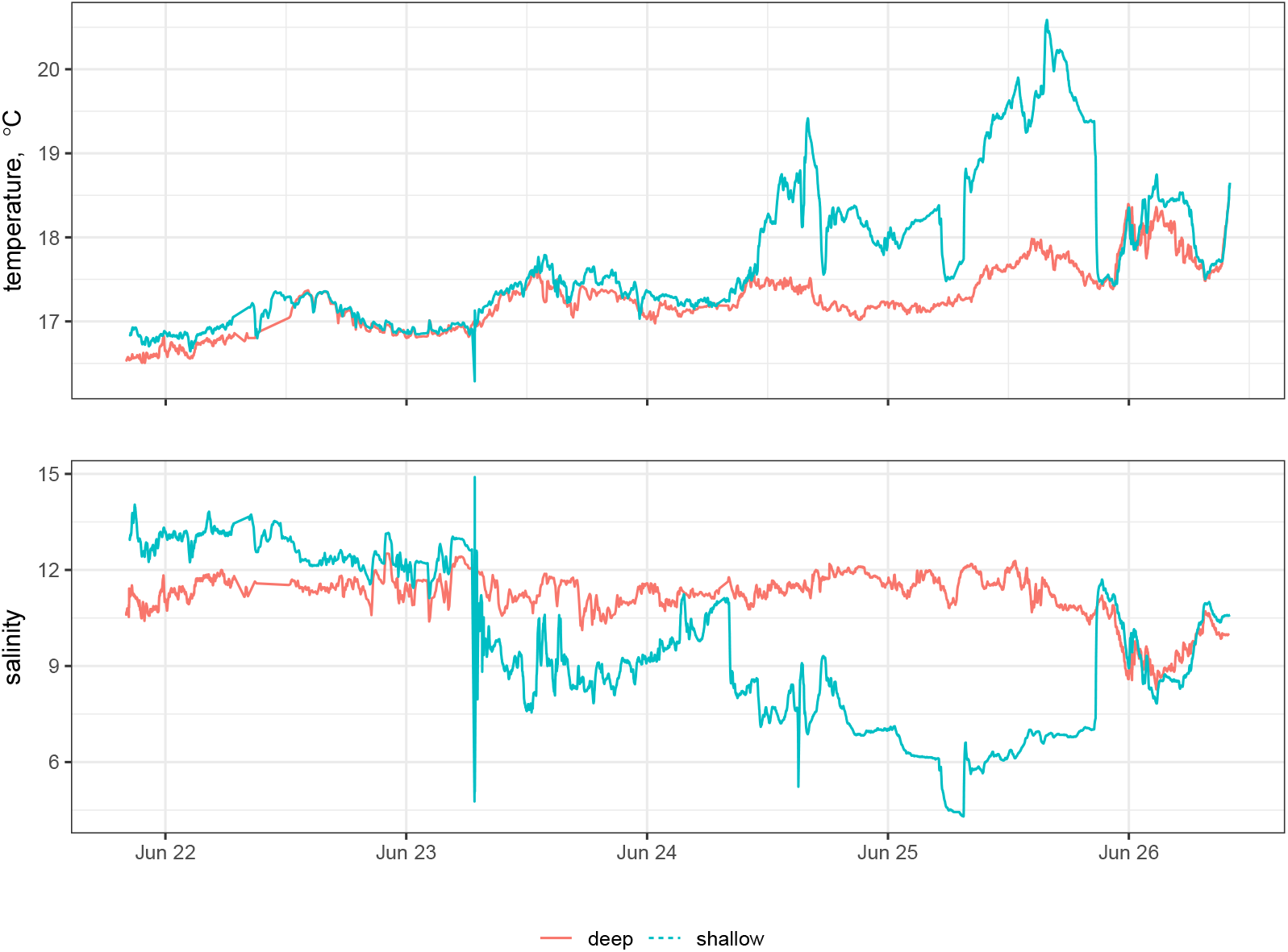
Seawater temperature and salinity during the experimental trial. Shallow refers to the seawater from the pump being set at different depths during the experiment. Deep refers to the pump fixed at 3m depth

